# Comparison of intradermal injection and epicutaneous laser microporation for antitumor vaccine delivery in a human skin explant model

**DOI:** 10.1101/861930

**Authors:** Sanne Duinkerken, Joyce Lübbers, Dorian Stolk, Jana Vree, Martino Ambrosini, Hakan Kalay, Yvette van Kooyk

## Abstract

Human skin is a prime vaccination site containing multiple antigen presenting cells (APC), including Langerhans cells (LC) and dendritic cells (DC). APC deliver antigens to lymph nodes for induction of adaptive immunity through stimulating antigen specific T- and B-cells. Since intradermal (ID) injections require specific training, easy applicable delivery systems like laser microporation are emerging. In mice, combination of laser treatment and allergen injection showed enhanced T-cell responses and skewing of B-cell responses without adjuvants. However, it remains to be elucidated whether laser microporation can alter human skin DC phenotype and function without adjuvants. In an *ex-vivo* human skin explant model, we compared ID injection and laser microporation as anti-tumor vaccination strategy. A melanoma specific synthetic long peptide and multivalent dendrimer with gp100 antigen were used as vaccine formulation to measure APC phenotype, emigration and ability to stimulate gp100 specific CD8^+^ T-cells. We show that skin APC phenotype and emigration capacity was similar after laser microporation and ID injection. However, laser microporation reduced vaccine uptake by APC, resulting in decreased induction of gp100 specific CD8^+^ T-cell activation. To conclude, in our human skin model ID injection remains the most potent strategy to deliver antigens to skin APC for T-cell induction.

**Graphical Abstract:** 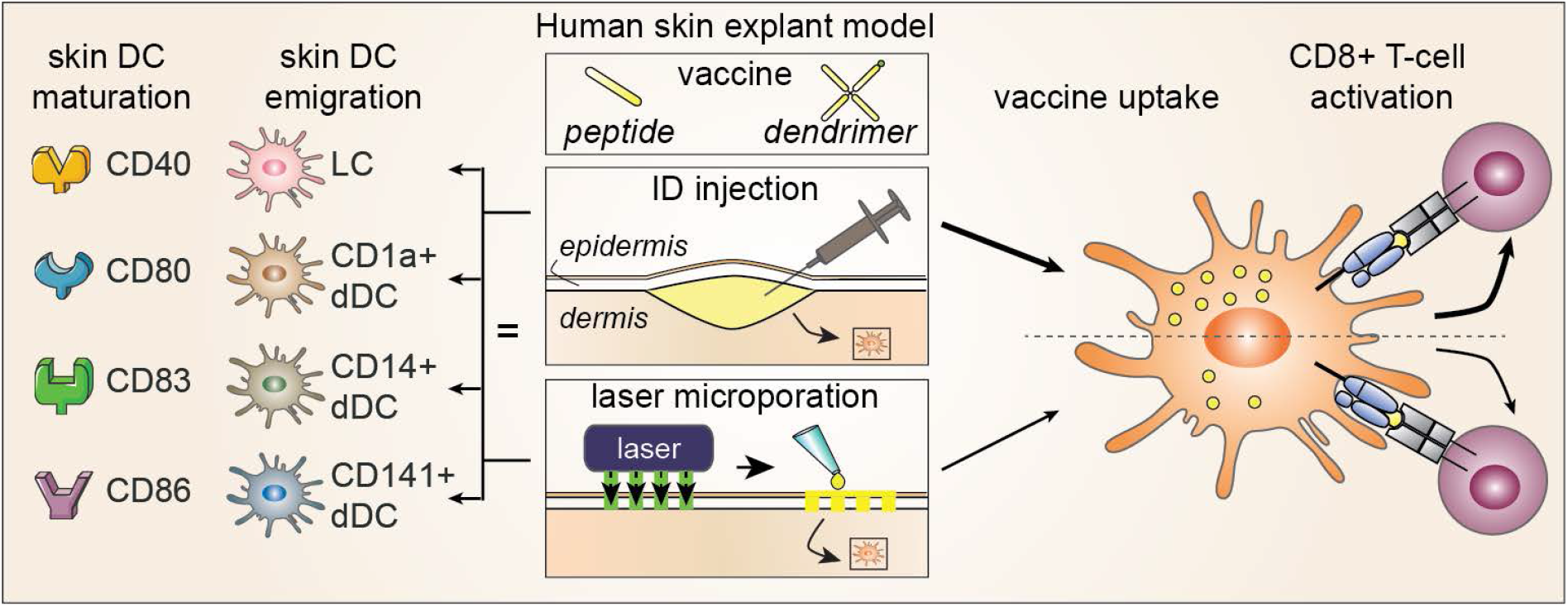

## INTRODUCTION

The human skin is a potent vaccination site as it contains numerous antigen presenting cells (APC), including epidermal Langerhans cells (LC), dermal dendritic cells (dDC) and monocyte derived macrophages (moMØ)^1^. APCs in the dermis can be subdivided into three distinct subsets: CD1a^+^ dDC, CD141^+^ dDC, CD14^+^ moMØ and a small population of Langerin^+^ dDC^1,2^. APC constantly scan their environment to detect changes in homeostasis, and are equipped with a plethora of pattern recognition receptors (PRR), such as toll-like receptors (TLR) that detect invading pathogens^3^. To help DC internalize antigenic content, DC express uptake receptors like C-type lectin receptors (CLR) that further process endocytosed pathogens into antigenic fragments to present on major histocompatibility complex (MHC) molecules. DC carry MHC presented antigenic peptides to draining lymph nodes for instruction of T-cell mediated adaptive immune responses^4^. LC and CD1a^+^ dDC are natural inducers of cytotoxic CD8^+^ T-cell responses, whereas the skin resident CD14^+^ moMØ can do so following specific CLR targeting and uptake^5^. CD141^+^ dDC can rapidly migrate to the skin draining lymphnodes and represent 20-40% of the DCs in the lymphnode ^1,2^. Furthermore, CD141^+^ dDC can skew the CD4^+^ T cell response to a th1 phenotype. Overall, the different skin APCs are excellent targets for vaccination strategies that can induce CD8^+^ T-cell responses required in the context of pathogens or tumors.

Compared to standard vaccine administration routes, skin immunization is more potent and dose-sparing^6^. Though, as intradermal (ID) injections require specific training and handling, most vaccination strategies still use subcutaneous or intramuscular vaccine delivery. In recent years new administration techniques have paved the way for easier and even pain free ID vaccination^7^. One of these techniques uses fractional laser ablation at a wavelength of water absorption to create clean micropores without thermally damaged ridges^8^. Depending on the settings, micropores can either disrupt the stratum corneum, reach into the epidermis or deeper into the dermis^9^. Multiple studies showed vaccine deposition within the skin already at minor disruption of the epidermis, thereby likely making vaccine particles available for endocytosis by skin APC^9^.

An important aspect for induction of adaptive immunity by DC, is the activation of PRRs which induces DC maturation and is often accomplished by adjuvant inclusion into the vaccine. Mature APC present antigens in MHC molecules, express co-stimulatory markers and secrete cytokines for proper T-cell activation in the lymph nodes^10,11^. The human skin is a plastic environment also containing non-immune cells such as keratinocytes and fibroblasts which can alter the environment by secretion of inflammatory cytokines^12^ and thereby activate skin resident APCs. Furthermore, dermal vasculature allows for influx of innate cells such as monocytes. Interestingly, recent murine studies showed immunomodulation upon nondestructive laser treatment in combination with ID vaccine injection. This included enhanced vaccine uptake, DC motility and increased CD4^+^ T-cell numbers and antigen specific antibody titers^13^. Since this technique still requires ID injection, other studies investigated the potency of destructive lasers creating micropores for vaccine delivery and enhancement of immune responses. Although laser microporation efficiently delivers vaccine particles into the dermis^14^, it was not directly compared to ID injection. Furthermore, the type of antigen appears to determine Th1 or Th2 skewing of immune responses and addition of adjuvant did not further enhance antibody titers compared to single laser treatment^15^. Thus, implying that laser microporation can be used for ID vaccine delivery and triggering of immune responses without a need for adjuvants.

Since both vaccination against pathogens and cancer require the induction of antibody responses and effective CD8^+^ T-cell responses, laser microporation may also be beneficial for anti-tumor vaccination strategies. A direct comparison between ID injection and laser microporation with tumor specific antigenic compounds needs to be tested in a human skin setting. In this study, we used a human skin explant model to investigate laser induced effects on human skin APC subsets. Using the P.L.E.A.S.E. laser device we compared epicutaneous laser microporation to ID injection of two tumor specific antigenic vehicles without adjuvant. Since tumor specific epitopes can be included in different types of vaccines, thereby favoring APC endocytosis^16^, we tested a linear synthetic long peptide (SLP) of the melanoma antigen gp100 and a larger multivalent dendrimer comprising multiple gp100-SLP bound to a scaffold. Both laser microporation and ID injection resulted in equal numbers human skin APC subset emigration with identical phenotype. Although laser microporation supplied both vaccine compounds to all human skin APC subsets, a lower amount accumulated compared to ID injection, that resulted in decreased gp100 specific CD8^+^ T-cell activation. Overall, our data shows that tumor specific compounds reach human skin APC more efficient through ID injection, resulting in efficient tumor specific CD8^+^ T-cell stimulation.

## RESULTS

### ID injection and laser microporation induce equal skin APC emigration

The skin consists of the epidermis covered by the stratum corneum (SC) and the dermis which are connected by a basal membrane containing tight-junctions that provide a barrier function^17^. To ensure proper vaccine delivery to APCs in the skin, both the SC and basal membrane need to be crossed. Laser microporation creates micropores in the skin with different depths depending on the laser settings. We verified that in a human skin explant model, laser ablation with 23.7J/cm^2^, four pulses and a depth of approximately 95μm as indicated by the P.L.E.A.S.E laser (**Fig.S1a**) was sufficient to penetrate through the epidermis and basal membrane, while leaving the dermis intact to allow vaccine delivery to LC, dDC and moMØ. Therefore, these settings were used for vaccine delivery in comparison to ID injection in a human skin explant model.

First, we evaluated whether laser microporation can enhance skin APC emigration compared to ID injection by analyzing subset ratio and absolute number of skin resident APC emigrated out of cultured skin biopsies (schematic representation **Fig.1a**). Using specific markers, MHC II^+^ (HLA-DR) human skin APC subsets were separated into the different epidermal and dermal subsets based on their marker expression (**Fig.S1b**), as previously described^18^. No differences were observed in skin APC subset emigration after two days of biopsy culture when skin was either untreated, ID injected or treated with laser microporation (**Fig.1b,c**). This indicates that laser microporation potentiates all human skin APC subsets to emigrate from the human skin equally well as ID injection.

**Figure 1.**
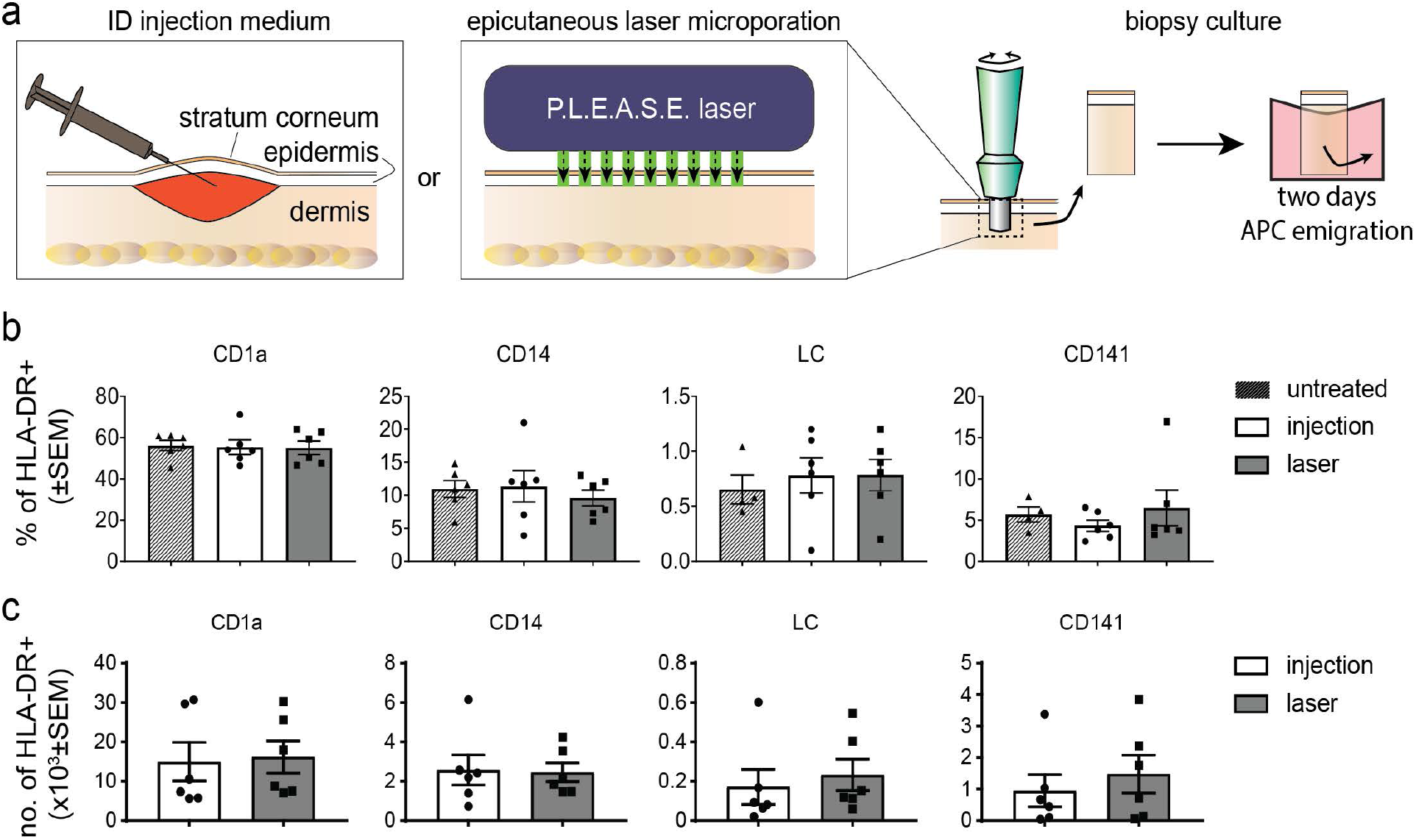
Equal human skin APC emigration following ID injection or laser microporation. (a) Human skin explants were untreated or treated with ID injection using insulin needles or laser microporation using the P.L.E.A.S.E. Professional laser device. Biopsies were cultured for two days followed by evaluation of emigrated skin APC subset ratio (b) and absolute number for ID injection or laser poration (c). n=6±SEM each dot represents a donor (Statistical analysis: Student’s t-test showed no significant differences).

### ID injection outperforms laser microporation for vaccine delivery to human skin APC

An important DC feature is their endocytic capacity for antigen or vaccine particle processing and subsequent presentation to T-cells in the lymph nodes. Antigen endocytosis by skin APC can be influenced by particle size and, as such, we used two sizes of synthetic vaccines to verify uptake by the different skin APC subsets following ID injection or laser microporation.

A single SLP containing the melanoma specific gp100-epitope for presentation in MHC I, was either injected as linear peptide (~3kD) or coupled to a dendrimer with four functional groups to create a larger multivalent peptide structure (~20kD). To evaluate uptake, the SLP amino acid sequence was elongated with an HA-tag and an AF488 was coupled to the dendrimer extremities (schematic representation **Fig.2a**). Using an insulin needle SLP or dendrimers were ID injected creating a small blister just under the epidermis of approximately 0.8cm^2^. For delivery following laser microporation the SLP or dendrimers were topically applied on an area of 1cm^2^. To avoid evaporation of the vaccine dissolvent, micropores were covered with Tegaderm followed by gentle massage of the applied vaccine area (schematic representation **Fig.2a**). Biopsies were cultured for two days to allow skin APC emigration. Following ID injection we could find HA-signal with FACS analysis in all APC subsets after two day emigration from skin biopsies (**Fig.2b,c**). Strikingly, application following laser microporation decreased uptake by all skin DC, moMØ and LC for both gp100-SLP (**Fig.2b,c**) and gp100-dendrimers (**Fig.2d,e**). No uptake for both gp100-SLP and gp100-dendrimers was found in the HLA-DR negative population (**Fig.S2a**). These data indicate that despite similar amount of vaccine molecule application, ID injection may facilitate better vaccine uptake by skin APC compared to topical application following laser microporation.

**Figure 2.**
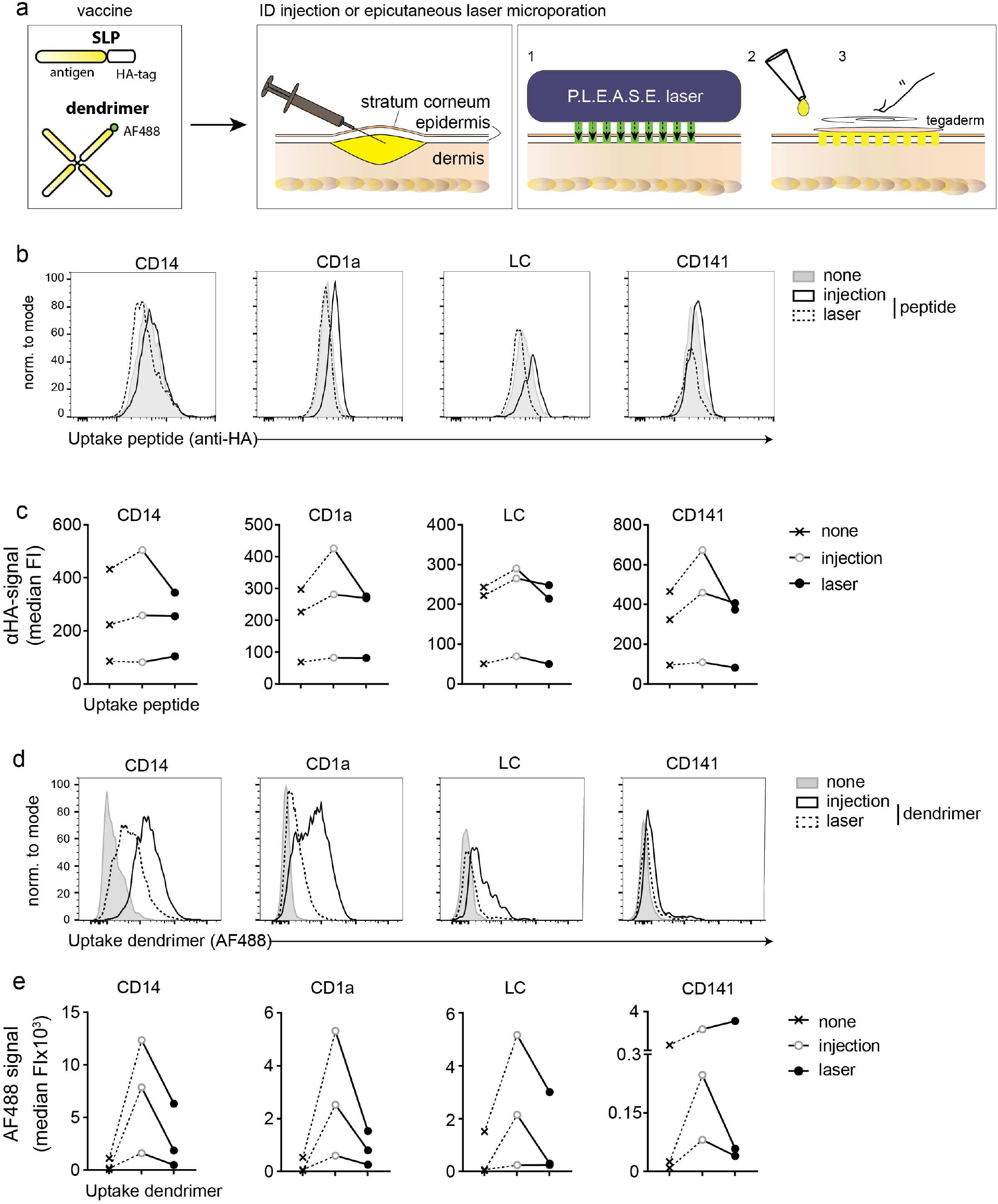
Superior vaccine uptake by skin APC following ID injection. (a) Human skin explants were used to evaluate in situ uptake of SLP or dendrimers by the different skin APC subsets following ID injection or laser microporation. (b,c) HA-signal of gp100-peptides compared to medium ID injection with the secondary antibody against the HA-tag (none) and (d,e) AF488 signal of multivalent gp100-dendrimers taken up by the different human skin APC subsets compared to medium ID injection (none). n=3 with representative histograms, each dot represents a donor. Symbols represent different treatments.

### Similar skin APC phenotype and skin micro-milieu following ID injection and laser microporation

Although skin APC subset emigration was similar following ID injection and laser microporation (**Fig.1b,c**) without an adjuvant, we wanted to verify if lowered antigen uptake by APC following laser microporation was due to phenotypical changes. Therefore, we analyzed co-stimulatory marker surface expression by the different skin APC subsets after two-day emigration. Overall, CD14^+^ moMØ showed lower expression levels of CD40, CD80, CD83 and CD86 compared to the other APC subsets, as previously described^19^. However, no differences in co-stimulatory marker expression after emigration were observed within the APC subsets between ID injection and laser microporation (**Fig.3**). Also, no differences were observed for CD83 and CD86 within the APC subsets between untreated and ID injected skin (**Fig.S2b**)

**Figure 3.**
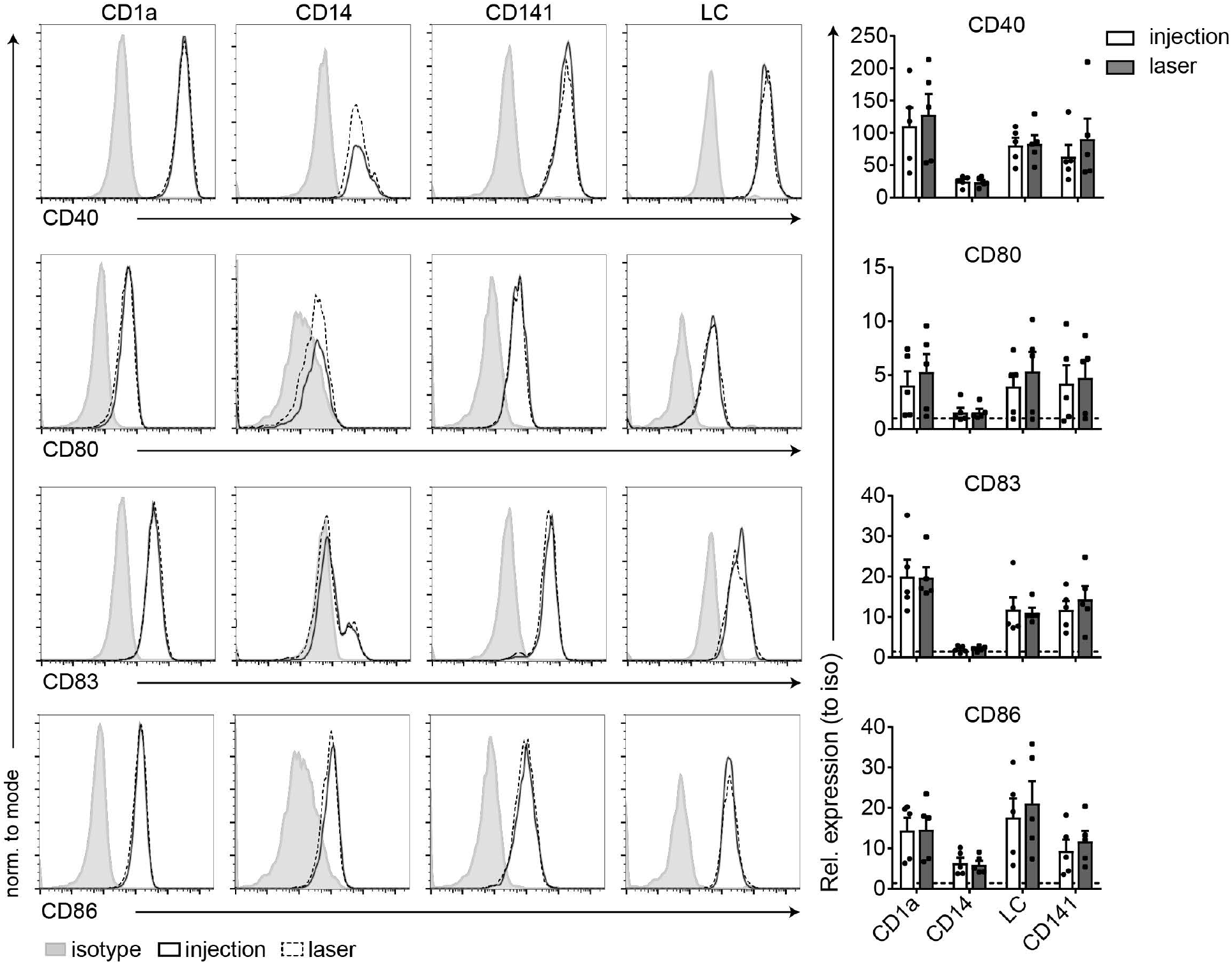
Similar co-stimulatory marker expression by skin APC subsets following ID injection and laser microporation. Human skin explants were either ID injected with culture medium or laser micro-porated followed by two-day biopsy culture. Emigrated skin APC subsets were analyzed for expression of maturation markers CD40, CD80, CD83 and CD86 using FACS analysis. n=6±SEM, each dot represents a donor. Representative histograms of one donor.

Human skin APC mature upon emigration^20^, so two days of emigration from the biopsy might overrule any initial differences in co-stimulatory marker expression induced upon laser treatment. Hence, we also elucidated cytokine secretion within the whole skin micro-milieu upon ID injection and laser microporation. During skin culture, biopsy conditioned supernatant was evaluated for presence of IL-1β, IL-6, IL-10, IL-12p70 and TNF-α at multiple time-points. Measurable levels of IL-6, IL-8 and IL-10 were detected after 6, 24 and 48 hour migration, whereas IL-1β, IL-12p70 and TNF-α were not detected. Figure 4 shows a significant decreased secretion of IL-6 and a trend for IL-8 after 48 hours when human skin was treated with laser microporation compared to ID injection (**Fig.4**). Overall, these data show that laser microporation does not induce phenotypical changes in APC marker expression and slightly alters cytokine micro milieu compared to ID injection in the absence of an adjuvant.

**Figure 4.**
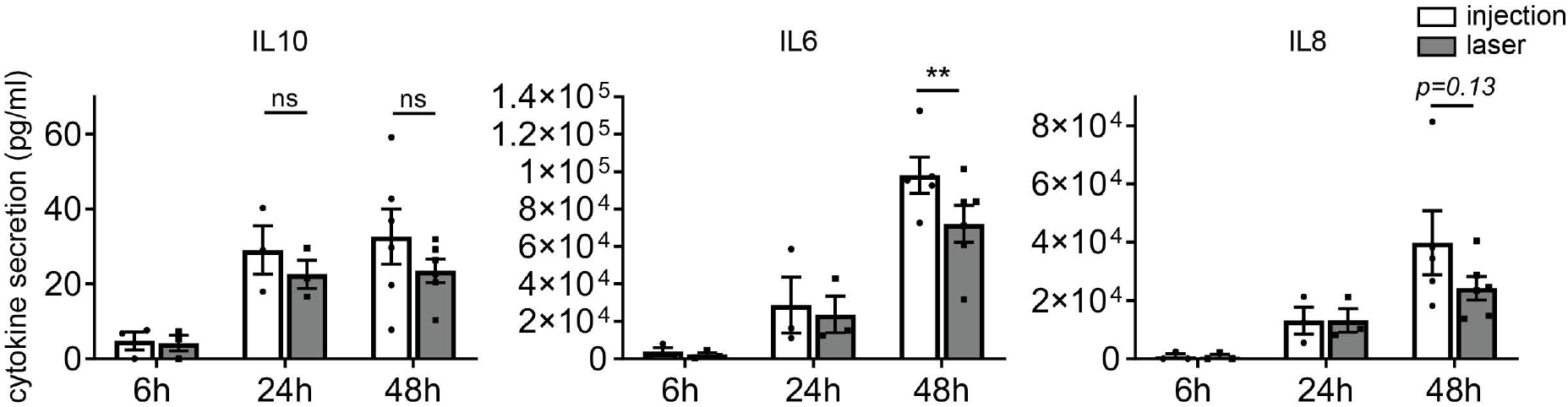
Skin micromilieu cytokine profile slightly changes upon laser microporation. Human skin explants were either ID injected with culture medium or laser micro-porated followed by two-day biopsy culture. Next, biopsy cultured supernatant was harvested and analyzed for secretion of cytokines after 6, 24 and 48 hours. Time course samples of 6h, 24h and 48h for 3 donors, 48h samples alone for 2 (IL-6) or 3 (IL-10 and IL-8) donors and each dot represents a donor.

### Higher CD8^+^ T-cell activation by skin APC following ID vaccine injection compared to laser microporation

Anti-tumor vaccination strategies rely on the induction of tumor specific CD8^+^ T-cells which requires APC to shuttle tumor antigens into the cross-presentation pathway for MHC I loading^21^. Therefore, cross-presentation of peptides and dendrimers by human skin APC was assessed following the different administration routes. We applied the gp100-SLP or the gp100-dendrimer as tumor vaccine to the human skin through injection or upon laser microporation. After two-day skin APC emigration, total skin APC were harvested and co-cultured with a gp100 specific T-cell clone recognizing the cross-presented vaccine gp100 minimal epitope in MHC I, and evaluated gp100 specific CD8^+^ T-cell activation by IFNγ secretion. Clearly, ID injected vaccines that targeted APC showed an increase in gp100 specific CD8^+^ T-cell activation for both the gp100-peptide and gp100-dendrimer vaccine (**Fig.5a,b** *open bars*). Surprisingly, but in line with the decreased uptake by skin APC following laser microporation (**Fig.2**), we observed that similar content of vaccine antigen delivered through laser microporation led to lower gp100 specific CD8^+^ T-cell activation compared to ID injection (**Fig.5a,b** *closed bars*).

**Figure 5.**
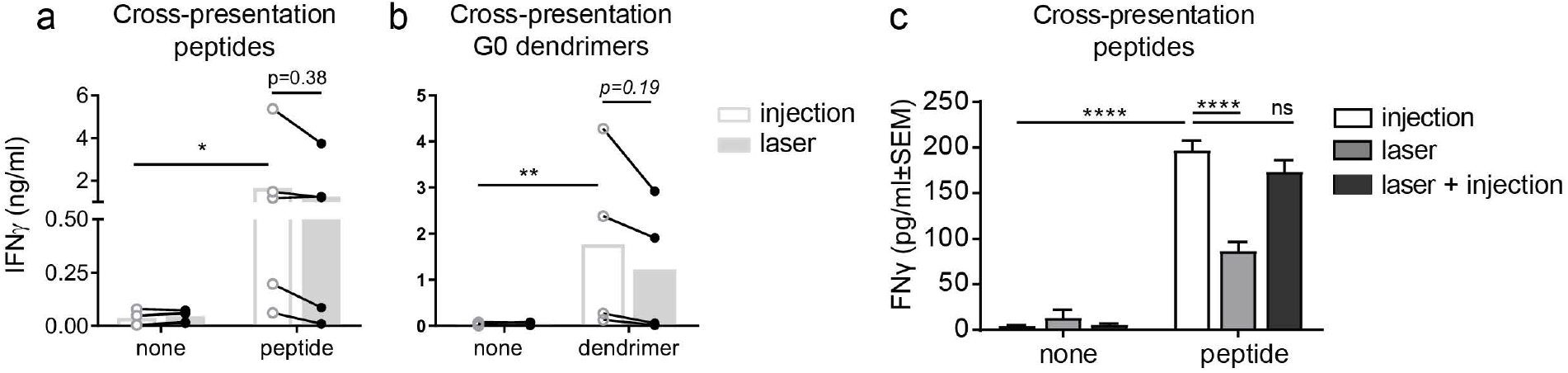
Higher gp100 specific CD8^+^ T-cell activation following ID vaccine injection. Melanoma specific gp100-peptides (a) or gp100-dendrimers (b) compared to a medium control (none) were either ID injected, applied topically following laser microporation or ID injected following laser microporation (c). After two-day biopsy culture emigrated APC were harvested and co-cultured with a gp100 specific T-cell clone. T-cell activation was measured by IFNγ secretion in the supernatant. (a) n=5, (b) n=4, each dot represents a donor. (c) representative of n=2 measured in triplicate.

To verify whether direct changes within the skin biopsy alter the capacity of skin APC to activate CD8^+^ T-cells, we first treated the skin with laser microporation followed by injection of the gp100-SLP vaccine in the skin. The capacity of the skin emigrated APC to activate CD8^+^ T-cells was restored as when the vaccine was only ID injected in the skin (**Fig. 5c**). This indicates that a decreased uptake of the vaccine by human skin APC following laser microporation results in reduced gp100 specific CD8^+^ T-cell activation.

## DISCUSSION

In this study we compared the use of ID anti-tumor vaccine delivery to human skin APC via injection or laser microporation without adjuvant. Laser microporation creating micropores with a depth just through the epidermis did not compromise human skin APC emigration or phenotype. However, tumor-specific vaccine particles were more efficiently delivered to the different human skin APC subsets upon ID injection compared to application following laser microporation. This resulted in higher tumor specific CD8^+^ T-cell activation following ID injection. Overall, our data show that in the ex-vivo human skin model, ID delivery of unmodified tumor-specific particles reaches human skin APC subsets best through ID injection.

For anti-tumor immune responses, the induction of cytotoxic CD8^+^ T-cells for tumor cell killing is important^22^. To achieve this, APC must shuttle exogenous derived tumor specific antigens into the cross-presentation pathway for loading on MHC I^21^. Cross-presentation by APC is influenced by mode of antigen uptake, but also signaling events occurring during antigen uptake. For example, concomitant TLR4 signaling enhances cross-presentation by retention of antigens in the endosomes through delay of endo-lysosmal fusion for antigen destruction^23^. As such, vaccines are often administered in the presence of TLR activating adjuvants to induce productive anti-tumor immune responses. However, laser microporation might induce so called adjuvanting effects e.g. by enhancing APC motility and antigen uptake. Our vaccination setting using unmodified peptidic compounds without adjuvant, did not benefit from laser microporation compared to ID injection. Leading to diminished activation of the CD8^+^ gp100 T cell clone, which appeared to be due to failure of particle uptake by the different APC subsets, although this might be underestimated due to processing of the SLP or dendrimer by the APC in the two-day culture and thereby removing the HA-tag or fluorescent label. Furthermore, there is a difference in diffusion of the products over time after laser treatment as shown by HX Nguyen et al ^24^ where the diffusion in vertical direction is highest in the first four hours and the horizontal diffusion in the skin after laser treatment is strongest after 8 hours. This will also impact uptake of the products by APCs over time.

The characteristics of vaccine compounds appear to dictate vaccine distribution and uptake by APC^14^. Interestingly, an important characteristic that can be altered to increase vaccine particle uptake is the size of the antigenic component^25^, however, even with the multivalent dendrimers we did not see a favorable contribution of laser microporation to uptake by skin resident APC. This is in line with previous studies where the molecular weight of particles did not influence vaccine distribution in the skin following laser microporation^14^.

Alternatively, the ability to specifically target vaccine formulations to APC via CLRs, such as DC-SIGN and the mannose receptor, which recognize specific carbohydrate structures, is an option^26^. Targeting vaccines can be created by coupling of carbohydrates to antigen specific vaccine particles. Using the grass pollen Phl p 5 protein functionalized with the mannose receptor binding carbohydrate mannan, Machado *et al* showed efficient targeting of skin APC^18^. Interestingly, this study showed synergy between targeting ability of the vaccine and laser microporation, where laser microporation enhanced uptake of the targeting particles in human skin APC compared to ID injection. In contrast, our study uses unmodified SLP and dendrimers without specific targeting characteristics that might be necessary for enhanced antigen delivery to human skin APC in combination with laser microporation.

In a murine study by Terhorst *et al*, XCR1 targeting anti-tumor vaccibodies were combined with (epi) cutaneous laser microporation and showed enhanced anti-tumor immune responses and both therapeutic and prophylactic efficacy^27^. Interestingly, here laser microporation induced immunomodulatory responses, thereby bypassing the need for adjuvant application. In our study we did not observe enhanced APC maturation as measured by co-stimulatory marker expression upon laser microporation. Though, co-stimulatory marker expression is often induced upon TLR activation. The immunomodulation of laser microporation does not appear to rely on signaling through TLR, but rather be an effect of signaling cascades initiated through local cell death and immune cell influx^27^. Unfortunately, we cannot study effects of laser microporation in the context of immune cell influx in the human skin explant model. Though, local cell death of KC is most likely induced which may alter local cytokine levels in the skin thereby affecting local (immune) cells and immune cell influx^28^. Interestingly, we observed a minor decrease in secretion of IL-6, IL-8 and IL-10 within the skin biopsy micro-milieu compared to ID injection. This hints to a micro-milieu with less attraction of e.g. neutrophils (IL-8). Importantly, this secretion of cytokines were measured within the total skin biopsy, whereas conflicting data is published when looking at the cytokine secretion profile of T-cells harvested from the spleen^8,29^. Where Weiss *et al* showed that deeper microporation into the dermis of mice increases Th1/2/17 cytokine secretion profiles by splenic T-cells^8^, Chen *et al* observed an overall decrease in cytokine responses^29^. Overall, it appears either immune cell influx induced upon laser microporation, or, skin layer dependent cell death is important for immunomodulatory effects of laser microporation.

Besides ablative fractional lasers (AFL), as used in this study, also non-ablative fractional lasers (NAFL) have been developed for the disruption of skin tissue without the formation of micropores. Combination of NAFL prior to ID injection enhanced uptake of unmodified vaccine through enhanced APC motility^13^. This resulted in both increased antigen uptake and migration toward the lymphatic system within the skin. Therefore, NAFL might be a better approach for intradermal vaccinations using unmodified antigenic compounds as used in our study, since we did not observe enhanced uptake following AFL treatment.

Overall, our study demonstrates that using the human skin explant model ID injection results in better antigen uptake by skin APC compared to laser microporation reflecting a superior tumor specific CD8^+^ T-cell activation.

## MATERIAL AND METHODS

### Human skin explant culture and DC isolation

Human skin explants were obtained within 24 hours after abdominal resection surgery (Bergman Clinics, Hilversum, The Netherlands) from healthy donors following informed consent as approved by the VUmc Medical Ethical Committee and research was performed according to the local guidelines and regulations of the Amsterdam UMC location VUmc. Obtained skin was rinsed with PBS substituted with 10μg/ml Gentamycin (Lonza) and subsequently ID injected using an insulin needle or porated using the P.L.E.A.S.E. Professional laser device (Pantec Biosolutions) with the following settings: fluency of 23.7 or 35,6 J/cm^2^, 4 pulses/pore, repetition rate 200Hz, pulse duration 75μs and 10% pore density. Per condition 8-12 biopsies were taken using 8mm biopsy-punches (Microtek, Zutphen, Netherlands) and cultured for 48 hours in 1 ml IMDM supplemented with 10% FCS, 10ug/ml gentamycin, penicillin/streptomycin (Lonza), and L-glutamin (Lonza) at 37°C and 5% CO2. Conditioned supernatant was collected at 6, 24 and 48 hours during culture for cytokine measurements. Following culture, skin biopsies were discarded and emigrated cells were harvested and pooled per condition for further analysis.

### Human skin section staining

For pore depth evaluation following laser microporation, skin biopsies were dried shortly on a tissue followed by emersion in TissueTek (Sakura Finetek, USA) and immediate freezing under liquid nitrogen. Skin biopsies were cut in 7μm sections (CryoStar NX70, ThermoFisher scientific), placed on 1% gelatin (Merck, Darmstadt, Germany) coated slides, dried and stored at −80°C. Prior to staining, sections were dried o/n at RT. Standard Haematoxylin Eosin (H&E) staining was performed and microscopic pictures were captured using the Leica DM6000 (Leica microsystems).

### Phenotypical analysis LC dDC and moMØ

Emigrated cells were washed in PBS with 1% BSA and 0.02% NaN3 and incubated for 30 minutes on 4°C with subset markers HLA-DR BV510 (clone G46-6, BD biosciences), CD1a APC (clone HI149, BD), CD14 AF700 (clone M5E2, Antibody Chain), CD141 BV711 (clone 1A4, BD), EpCAM BV421(clone 9C4, Antibody Chain) and maturation markers CD86 FITC (clone BU63, ImmunoTools), CD80 FITC (clone 2D10, Biolegend), CD40 PE-Cy7 (clone 5C3, Biolegend), CD83 PE-Cy7 (clone HB15e, BD) and Fixable Viability Dye eFluor780 (eBioscience). Cells were measured by flow cytometry (Fortessa X-20, Beckton Dickinson, San Jose, CA, USA) and analyzed with FlowJo (V10, Tree Star, Ashland, OR, USA).

### Cytokine secretion skin biopsies

Cytokine secretion was measured using sandwich ELISA according to manufacturer’s protocol with specific cytokine antibody pairs for IL-1β, IL-6, IL-8 (BioSource), IL-10, IL-12p70 and TNF-α (eBioscience).

### Peptide and dendrimer synthesis

In short, the gp100-SLP containing the CD8^+^ T-cell epitope (bold) and HA-tag (underlined) (Thz-VTHT**YLEPGPVTA**NRQLYPEWTEAQRLD<u>YPYDVPDYA</u>-C) was synthesized by microwave assisted solid phase peptide synthesis using Fmoc chemistry on the Liberty blue peptide synthesizer (CEM).

A generation 0.0 PAMAM dendrimer core was functionalized with maleimide followed by coupling at the C-terminal end of the gp100-SLP containing a specific linker (Thz-VTHTYLEPGPVTANRQLYPEWTEAQRLD-**(Abu)_3_**-C). Next, AF488-maleimide was added at the unmasked N-terminal (Thz) end for tracking. Mass and purity of all steps were confirmed by UHPLC-MS (Ultimate 3000 UHPLC, Thermo Fisher) hyphenated with a LCQ-Deca XP Iontrap ESI mass spectrometer (Thermo Finnigan) using a RSLC 120 C18 Acclaim 2.2um 2.1 x 250 mm column. Mass spectrometer analysis was measured in positive mode (**Fig.S3**).

### Human skin APC *in-situ* vaccine uptake

HA-tagged peptides or AF488-labeled dendrimers were diluted in serum-free IMDM and 20μl was ID injected or 10μl (2x concentration) was applied topically on 10mm^2^ following laser microporation and gently massaged (10seconds). Laser microporated areas were covered with Tegaderm film (Newpharma, Luik, Belgium) to avoid evaporation. Ten 8mm punch-biopsies were taken per condition and cultured as described above. For peptide uptake, APCs were fixed and permeabilized using a Cytofix/Cytoperm kit (BD) according to manufacturer’s instructions and stained with an anti-HA-AF488 (clone 6E2; Cell Signaling) antibody for 30minutes at 4°C. Cells were stained for subset markers and analyzed by flow cytometry as described above.

### Human skin APC antigen presentation

For antigen presentation a specific T-cell clone recognizing the gp100 HLA-A2 minimal epitope was produced and cultured as previously described^30^. HLA-A2^+^ skin was used for vaccine injection or application after laser microporation and 8 biopsies were taken and cultured as described above. Harvested cells were co-cultured in a 96-well round bottom plate with the T-cells in a 1:5 ratio (triplicate). After 24 hours supernatants were harvested and T-cell activation measured using an IFNγ Ready-Set-Go kit (eBioscience) according to manufacturer’s protocol.

### Statistical analysis

GraphPad Prism (V7.02) was used for statistical analysis. Student’s T-test was used to compare percentage or number of cells. For group comparison two-way ANOVA followed by Tukey multiple comparison test were used with paired analyses for multiple donors. P-values <0.05 were considered to be significant.

## Supporting information

Review report

## DATA AVAILABILITY

No datasets were generated or analyzed during the current study.

## AUTHOR CONTRIBUTIONS

S.D., J.L., Y.v.K. Conceptualization; Y.v.K. Funding Acquisition; S.D., J.L., D.S., J.V. Investigation; S.D., J.L. Methodology; S.D., Y.v.K. Project Administration; M.A., H.K. Resources; Y.v.K. Supervision; S.D., J.L., D.S. Validation; S.D. Visualization; S.D., Y.v.K. Writing - Original Draft Preparation; all authors Writing - Review and Editing

## ACKNOWLEDGEMENTS

This work was funded by the European Research Council (ERC Advanced 339977-Glycotreat S.D, J.L, Y.v.K) and the Dutch Cancer Society (VU2014-7200 D.S.). We would like to thank Pantec Biosolutions for use of their ablative P.L.E.A.S.E. laser device and Bergmanclinics for healthy donor skin supply to conduct this research. Many thanks to the GlycO2peptide, O2Flow Cytometry and AO2M Microscopy facilities of Amsterdam UMC (VUmc) for construct synthesis and technical support.

