## Supplementary material for "Comparison of intradermal injection and epicutaneous laser microporation for antitumor vaccine delivery in a human skin explant model": Review report

Reviewer comments:

Reviewer #1 (Remarks to the Author):

Duinkerken et al performed an interesting study to compare laser-assisted skin vaccination versus established intradermal application. They conclude from their data that intradermal vaccination is superior to the laser-assisted approach, however, this conclusion is not supported by their data. Please see the specific concerns listed below to illustrate this. Moreover, there are some discrepancies, e.g. the uptake of vaccines after laser-formation of pores is poor (Fig 2), however, the T cell responses induced are pretty good (Fig5). In many figures experiments with more donors would have been helpful to get to a convincing conclusion, as results with human skin can vary a lot depending on the UV-exposure, body site etc.

*Response: We would like to thank the reviewer for the time invested in reading and commenting on this manuscript. In the following point by point reply we address all the comments and remarks of the reviewer. One of the common remarks is the small number of donors per experiment figure. We would like to point out that this article is providing negative data that the laser poration is not better then dermal injection in this steady state human dermal skin model and therefore, we show the limited number of donors. For some of the experimental figures we have added in a number of donors as indicated in the point by point reply.*

There are several points that need to be addressed:
- How did the authors measure the penetration depth (e.g., 95 micrometers) of the laser? Or was this the value that was calculated and indicated by the laser device? Please specify.
*Response: the micropore depth is indicated by laser device itself. This information is now included in the manuscript page 5 line 91-92*

- Fig.1c: There are more CD141+ than LCs although the LC are more abundant in the skin. Do you have an explanation why there are so few LCs found in the emigrated DC?
*Response: In previous research we have always observed the same amounts of LC and CD141$^+$ cells that emigrate from the skin without adjuvants like GM-CSF/IL-4 or LPS (C. Fehres et al, European Journal of Immunology, 2014/ Y. Machado et al, Journal of controlled release, 2017). This was also reported by Chung-Ching Chu et al (JEM, 2012) for CD141+ cells and by Dinja Oosterhoff et al (The journal of Immunology, 2013) for LCs.*

***In this paper you show 10-times more CD141+DC than LC, so this is far away from same amounts. And the paper by Frank Nestle's group did not compare LC and CD141+DC as well as the other paper from Tanja de Gruijl's lab does not even investigate CD141+DC in their skin explant cultures. So why?***

- Fig.2: Antigens are not taken up at all after laser treatment. However, the migratory DC can induce T cell responses as shown in Fig.5. How does this fit together?
*Response: Indeed, the signal of antigen uptake is clearly decreased following laser microporation. Especially the signal for uptake of the SLP is abrogated following laser microporation. We think this is most likely due to the use of an HA-tag which might be affected by the processing during the two-day DC emigration, thereby underestimating the amount of antigen uptake. Though, oppositely to SLP, for dendrimers there is still a clear signal of AF488 (fig.2d), especially in CD14$^+$ and CD1a$^+$ dDC, albeit lower following laser microporation compared to injection. Also, here the signal Is measured*

*following two-day DC emigration, giving possible underestimation of total antigen uptake due to processing. Since we do find antigen presentation following two-day emigration it is apparent that the antigenic content from both SLP and dendrimer vaccines are sufficient for the induction of T cell responses, within the two-day time-frame. We concluded therefore that the read-out for antigen presentation is more sensitive than intracellular detection by HA, and fluorescence. We included this in the discussion on page 10 line 189-191)*

- Fig. 2: For control purposes and to put the data into a better perspective, the authors should also show the (absent?) uptake of SLP or dendrimers into the non-APCs, i.e., the HLA-DR-negative (or CD45-negative) cells.
<u>Response:</u> *We now included the control uptake by non-APCs, defined as HLA-DR- population, in the manuscript at page 7 lines 123-124 and in supplementary figure 2*

***There is uptake of vaccines in HLA-DR-negative cells, this does not fit to your statement. This data set shows that there is a lot of unspecific uptake of vaccines into non-immune cells, which is an important observation that needs to be discussed.***

- Fig.2: From a vaccination point-of-view it would be interesting to see, how well the SLPs/dendrimers are retained in the skin APCs. The authors should compare the 48h migratory cells with freshly isolated whole skin suspensions containing the same subsets of APCs. Are fluorescence intensities higher shortly after application of the antigens?
<u>Response:</u> *We agree with the reviewer that it is interesting to see whether the SLPs/dendrimers are retained in the skin APCs. However, this is difficult to compare between short (4 hours) and 48 hours incubation as our SLPs/dendrimers are both taken up and processed by the APCs and thereby the HA/fluorescent tag is removed due to the processing. At short 4 hours interval there is mostly uptake and little processing whereas at the 24 hours timepoint there is both uptake and processing. Furthermore, there is a difference in diffusion of the products over time after laser treatment as shown by HX Nguyen et al (International Journal of Pharmaceutics, 2018) where the diffusion in vertical direction is highest in the first four hours and the horizontal diffusion in the skin after laser treatment is strongest after 8 hours. This will also impact uptake of the products by APCs over time. We included this the discussion page 10 lines 191-195.*

- Fig.3: What are the grey histograms? Isotype controls or non-treated skin explant? As the authors already mentioned emigrated skin DC are activated so it is not surprising that maturation markers are comparable between both treatments. It would have helped to show the emigrated DC from untreated control skin in comparison.

<u>Response:</u> *Grey histograms represent isotype controls as depicted in the figure. In the past we have not observed differences between emigrated DCs of non-treated (or non-injected) versus medium injected skin biopsies. Since there is limited amount of skin available per experiment, we decided to use medium injection or topical application following laser microporation to evaluate the effect of tissue handling, with an emphasis on the effect of laser microporation, on human skin DCs in the absence of the vaccine. For clarification purposes we included in the manuscript on page 7 lines 134-136 and supplementary figure 2b the comparison between untreated and ID injected skin for the migration markers CD83 and CD86.*

To investigate phenotypic changes in skin DC by treatments, the authors need to perform whole skin digestion after few hours and 1-2 days to investigate differences in DC phenotype. Alternatively, microscopy of skin sections at different time points after treatment and skin explant culture should show differences in DC activation.
*Response: We agree it would be interesting to evaluate differences in DC phenotype within a few hours, compared to a few days. However, enzymes needed for whole skin digestion can affect extracellular marker expression by cleavage of such receptors (Botting et al. 2017 JLB), thereby possibly introducing a bias in receptor expression. Furthermore, induction of maturation also induces migration of skin DCs as described by D. Oosterhoff et al (Journal of Immunology, 2013), which will thus no longer be available in the whole skin tissue following 1-2day culture. As such, results might be skewed since "left behind" skin DCs might actually be the ones not affected by the different treatments and therefor have equal phenotypes as DCs harvested shortly following treatment.*

- Fig. 4: The authors claim that cells in laser treated skin secreted less cytokines than in intradermal injected skin. The results shown in the figure are not statistically significant and not all data points are shown.
The figure legend states that 4 donors are shown, but in some graphs only two dots are visible. So, this conclusion is an overinterpretation of the data. Human donors can vary a lot, thus I would highly recommend to perform additional experiments and include more donors. In addition, it is possible to measure cytokines in tissue protein lysates, which would have given a better picture of which cytokines are produced in the treated skin.

*Response: We agree with the reviewer that some data points were missing from the graph and that we should include more donors, therefore, we have measured the cytokines in 3 more donors at the 48 hours' time point and added these in the graphs as depicted in figure 4. With the addition of these 3 donors, the decrease in cytokine expression with laser poration compared to ID injection showed a trend towards IL-8 secretion and a significant decrease for IL-6 at 48 hours. This was adjusted at page 8 lines 145-147. Figure 4 was adjusted with the right number of donors.*

- Fig.5: Again, the conclusion here is a not really supported by the data. In Fig.5a statistical significance is not reached, again it would have been helpful to include more donors. And as mentioned above it is surprising to see a T cell response when the vaccines after laser pretreatment are not taken up by migratory DC.
*Response: In figure 5a and 5b the number of donors is increased to N=5 and N=4, respectively. Still the difference between ID injection and laser treatment is not significant, but revealed a clear trend in decreased cross presenting both the gp100-SLP and gp100-dendrimers after laser poration. Nevertheless, the uptake of mostly the SLP is almost abrogated and we would expect a bigger difference in cross presentation. As mentioned above in Fig 1C, we think the use of an HA-tag for the SLP gives an underestimation of antigen uptake following two-day emigration due to processing of the peptide and thereby the HA-tag within this time-frame. Therefore, we see some cross presentation of the gp100-SLP and -dendrimer.*

***Please rephrase this in the results and mention that these results are not statistically significant, only a trend!***

-In Fig.5c it is not clear what representative of 2 donors (which is again a low number!) means, doing statistics on experimental triplicates is not helpful. To prove that the combi works as well as injection more donors need to be included.
<u>Response:</u> *We agree that inclusion of more donors would be helpful. Though, the main message of fig.5c is not to prove that combination of injection and laser microporation works. Here we wanted to verify whether the decreased antigen presentation observed by laser microporation was a result of the decreased antigen uptake due to e.g. vaccine distribution, or, due to changes in skin micromilieu or DC phenotype following laser treatment. Since we see restoration of AGP when the vaccine is injected following laser poration, we think that the differences in vaccine distribution for uptake by DC is suboptimal upon laser microporation and not due to phenotypical changes. We highlighted in the text that relatively low numbers of donors are tested. However, in the n=2 the standard deviations are very low of the triplicates.*

***These data are important for the paper as it tries to give an explanation for the impaired antigen cross-presentation after laser microporation. I am still not convinced by the way the data is displayed in fig5c. that shows statistics of triplicates from one representative experiment (=1 donor). The authors should show both donors and present these results in a more descriptive way as no clear conclusion can be made from 2 donors. Or repeat the experiment with more donors.***

General comments:
- The population of CD14+myeloid cells that was formerly called CD14+dermal DC was shown to be a transient monocyte-derived macrophage population (MCGovern N et al, Immunity 2015). The authors should explain why they are still using this terminology
<u>Response:</u> *The reviewer addresses an important point, and we adjusted the terminology for CD14+ dermal APCs to CD14+ moMØ in the whole document. We also point out in the introduction that these CD14+ moMØ are resident MØ in healthy skin at page 3 line 36.*

- Please indicate in all figure legends of data points correspond to donor, missing in Fig.2

<u>Response:</u> *We added this to the figure legend of figure 2.*

Reviewer #2 (Technical Comments to the Author):

The present manuscript investigates anti-tumor vaccine delivery in human skin. The experimental approach compares a classical intradermal injection and a laser microporation followed by topical application of the vaccine. The vaccine administrations are performed in a human skin explant model and the study is focused on four DC subsets which emigrate from skin biopsies in a two-day culture. The authors conclude that the ID injection results in better antigen uptake by skin DC compared to laser microporation.

Response: *We would like to thank the reviewer for the positive feedback and time invested in reading and commenting on this manuscript. One of the common remarks is the missing comparison to the untreated skin. This was added in multiple figures and supplementary figures as stated in the rest of this point by point reply.*

Major concerns

1. The authors show that ID injection and laser microporation induce similar effects on skin DC emigration. Nevertheless, they do not show the spontaneous DC emigration from untreated skin. The results presented do not show whether the DC emigration is the result of vaccination or culture condition.

Response: *We agree with the reviewer that the spontaneous emigration is an important comparison. Therefore, the spontaneous DC emigration (untreated skin) per subset as a percentage of HLA-DR+ cells is now included to Figure 1b in the manuscript and also mentioned in the text at page 6 line 102.*

***And why not not in fig1c? numbers are quickly calculated!***

2. Figure 1b and 1c show the same results expressed differently. The total number of cells (Figure 1c) is more appropriate but it should be done more accurately by calculating the absolute cell number per gram of tissue.

Response: *Figure 1b and 1c show indeed the same results, because we think it is clearer to show both the percentages and number of cells, so, other readers could compare the cell percentage to other articles as well. We don't agree that the calculated cell number is more accurately per gram of tissue in our setting, as these tissue biopsies are floating in medium and also taking up medium. This would bias the calculation in cells per gram of tissue.*

***You could show it as numbers per biopsy!***

3. The authors show the expression of activation markers on the DC subsets after each vaccination method. Similarly to concern number 1, the expression of activation markers should be compared to DC spontaneously emigrated from the unvaccinated biopsies.

Response: *In the past we have not observed differences between emigrated DCs of non-treated (or non-injected) versus medium injected skin biopsies. Since there is limited amount of skin available per experiment, we decided to use medium injection or topical application following laser microporation to evaluate the effect of tissue handling, with an emphasis on the effect of laser microporation, on human skin DCs in the absence of the vaccine. For clarification purposes we added in the comparison between untreated and ID injected skin for the migration markers CD83 and CD86 in the manuscript on page 7 lines 123-124 and supplementary Figure 2b.*

4. The authors show the production of three cytokines out of six measured in the culture medium. Similarly to concerns number 1 and 3, the unvaccinated skin group is missing. The authors claim (p10) that "laser microporation … slightly alters cytokine micro milieu compared to ID injection…". However, it would be of greater interest to show the effect of each vaccination method in comparison to the cytokine production in untreated skin. =

*<u>Response:</u> As mentioned above, there is limited skin available per experiment and therefore, we decided to use medium injection or topical application following laser microporation to evaluate the effect of tissue handling. As such we do not have the possibility to add the cytokine production of untreated skin to the comparison.*

5. The authors claim that CD8+ T-cell activation by skin DC is higher after ID injection than after laser microporation. However, the results were obtained with an antigen-specific T cell clone which activation is directly dependent on the amount of antigen presented and it reflects the amount of antigen loaded in the DC. In an anti-tumor vaccine context, it would be of greater interest to show that the antigen-loaded DC are able to stimulate naïve CD8+ T cells.

*<u>Response:</u> The reviewer is indeed correct when stating the activation of the CD8+ T cell clone is a measure of DC cross-presentation and subsequent antigen loading on MHC-I. Unfortunately, we cannot evaluate activation of naïve CD8+ T cells since the gp100 antigen used in our vaccines is not naturally occurring in healthy human blood T cells. Though, we think the assay used in this study is useful to evaluate whether human skin DCs can efficiently harbor the vaccine following different delivery methods in situ, for subsequent processing and presentation. We highlight the use of the CD8+ gp100 T cell clone in the discussion at page 10 line 188-189.*

Figure 5 should be included in Figure 2: both figures concern the amount of antigen loaded in the skin DC.

*<u>Response:</u> We disagree with the reviewer that figure 5 should be combined with figure 2, as figure 5 shows the ability of the APCs to activate a T cell clone by cross presenting an antigen instead of taking up an amount of antigen by the moDCs. Moreover the order of the story is than completely changed with no proper ending.*

Minor concerns
1. The role of CD141+ DC should be mentioned in the introduction.

<u>Response</u>*: We now added in the following text in the introduction (page 3, line 37):* $CD141^+$ dDC can rapidly migrate to the skin draining lymphnodes and represent 20-40% of the DCs in the lymphnode (s. Kashem, Annual review of immunology, 2017). Furthermore, $CD141^+$ dDC can skew the $CD4^+$ T cell response to a th1 phenotype.

2. FigS1a should show a scale bar.

*<u>Response:</u> We adjusted the figure with a scale bar.*

3. "none" condition in Figure 2 and Figure 5 should be explained in the figure legends.

*Response:* *We added the explanation for the "none" condition in the figure legends.*

4. In the Material and Methods section, the phenotypical analysis of skin DC by cytometry does not mention any procedure to block the Fc receptors expressed by DC. If this is not a manuscript omission, the authors should show that it does not jeopardize the accuracy of the analysis.

*Response:* *In the manuscript we included the isotype controls to show an estimate of the non-specific binding of the FAB part of the antibody as this could be Fc receptor mediated or non-specific.*
