## Supplementary material for "Comparison of intradermal injection and epicutaneous laser microporation for antitumor vaccine delivery in a human skin explant model": Fig.S

### Supplementary figures

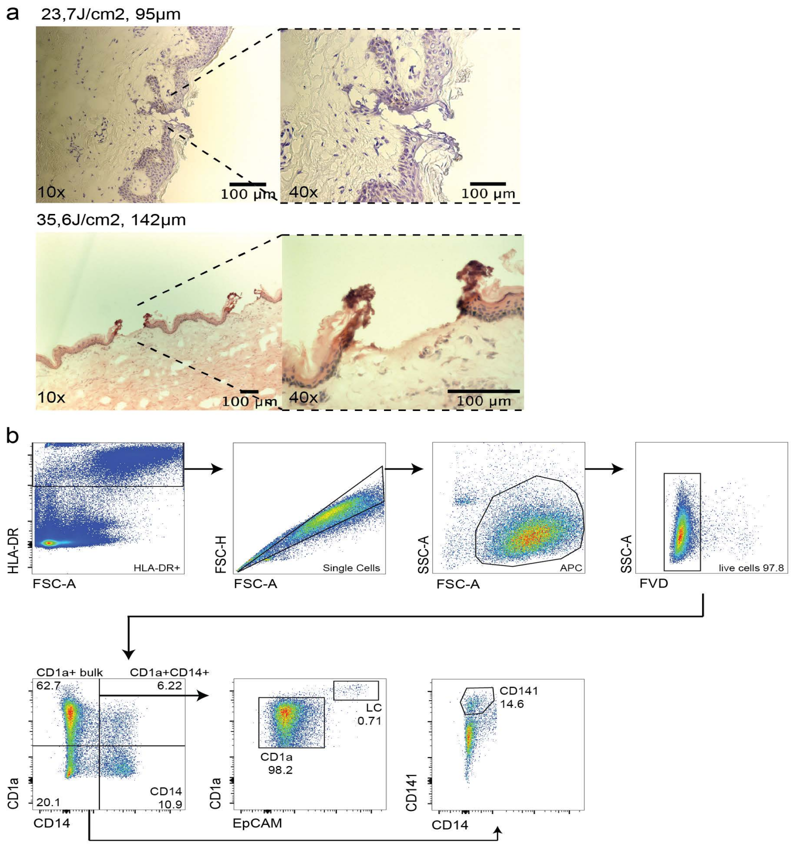


**Supplementary figure S1** (a) H&E staining of laser micro-porated human skin using 4 pulses and 23,7J/cm2 (top) or 35,6J/cm2 (bottom). (b) Gating strategy of human skin DC subsets following two day emigration using FACS.

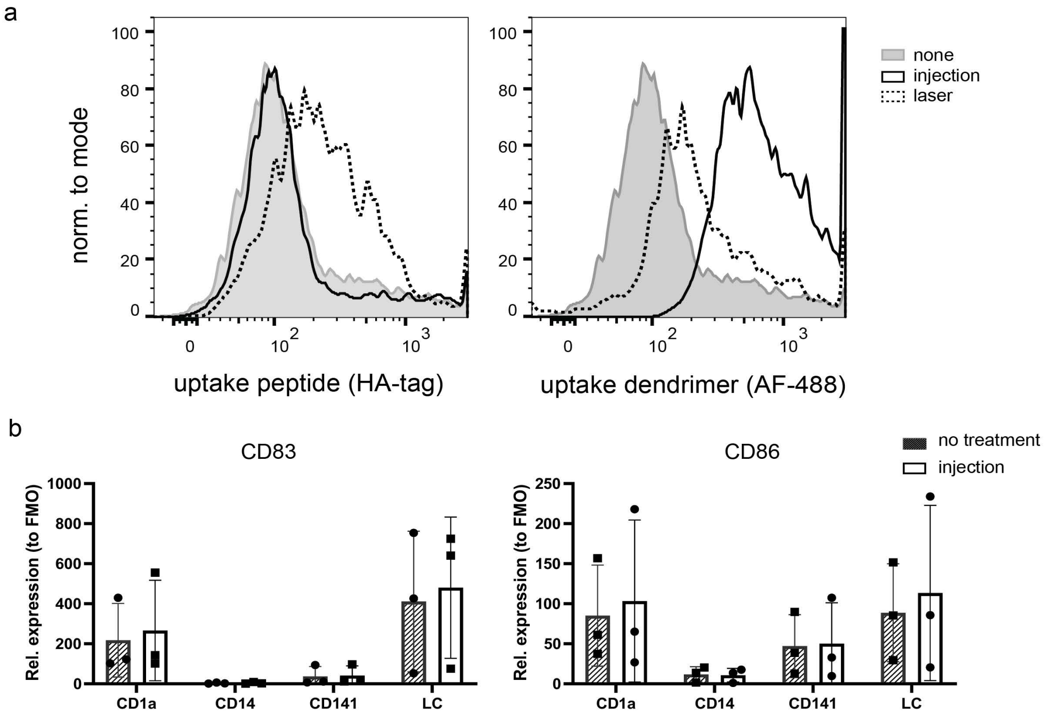


**Supplementary Figure S2** (a) Uptake of the SLP peptide with HA-Tag on the left and dendrimer with AF-488 on the right in the HLA-DR- population for medium treated (none), ID injection and laser poration. (b) CD83 and CD86 relative expression to FMO in all skin APC subsets for untreated and medium ID injections.

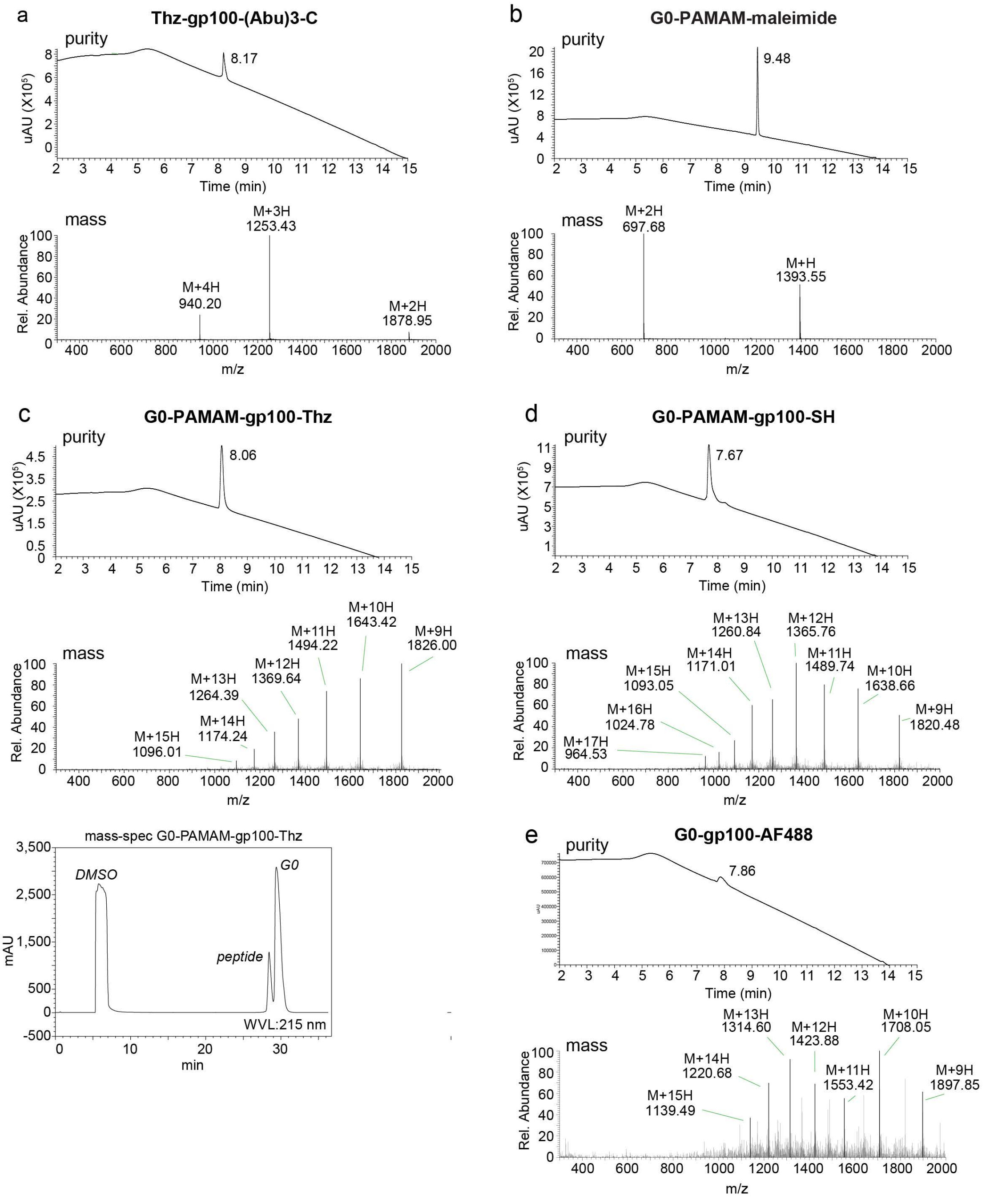


**Supplementary figure S2** Analysis of peptide and dendrimer synthesis Purity and mass as measured by UHPLC-MS and mass spectrometer system of SLP coupled to dendrimers (a) and G0 PAMAM dendrimer core functionalized with maleimide (b). coupled to SLP (c), N-terminal unmasking (d) and AF488 coupling (e).
